# A pH-sensitive switch activates virulence in *Salmonella*

**DOI:** 10.1101/2022.12.15.520350

**Authors:** Dasvit Shetty, Linda J. Kenney

**Author notes:** To whom correspondence should be sent.

## Abstract

The transcriptional regulator SsrB acts as a switch between virulent and biofilm lifestyles of non-typhoidal *Salmonella enterica* serovar Typhimurium. During infection, phosphorylated SsrB activates genes on *Salmonella* Pathogenicity Island-2 (SPI-2) essential for survival and replication within the macrophage. Low pH inside the vacuole is a key inducer of expression and SsrB activation. Previous studies demonstrated an increase in SsrB protein levels and DNA-binding affinity at low pH; the molecular basis was unknown (Liew et al., 2019). This study elucidates its underlying mechanism and *in vivo* significance. Employing single-molecule and transcriptional assays, we report that the SsrB DNA binding domain alone (SsrBc) is insufficient to induce acid pH sensitivity. Instead, His12, a conserved residue in the receiver domain, confers pH sensitivity to SsrB allosterically. Acid-dependent DNA binding was highly cooperative, suggesting a new configuration of SsrB oligomers at SPI-2-dependent promoters. His12 plays a role in SsrB phosphorylation; substituting His12 reduced phosphorylation at neutral pH and abolished pH-dependent differences. Failure to flip the switch in SsrB renders *Salmonella* avirulent and represents a potential means of controlling virulence.

## INTRODUCTION

*Salmonella enterica* serovar Typhimurium is a pathogen that causes gastroenteritis in humans and a typhoid-like disease in the mouse. *Salmonella* pathogenicity is largely conferred by the presence of horizontally-acquired virulence genes encoded within genomic regions called *Salmonella* pathogenicity islands (SPIs) (Hensel, 2000; C. A. Lee et al., 1992). The most well-characterized genomic islands are SPI-1 and SPI-2, which encode two distinct type three secretion systems (T3SS), as well as genes encoding secreted effectors that are important for pathogenesis. The SPI-1 T3SS aids in the initial attachment and invasion of the intestinal epithelium (Galán & Curtiss, 1989; Mills et al., 1995), while SPI-2 genes play an essential role in the survival of *Salmonella* within the macrophage vacuole and its subsequent maturation into a *Salmonella*-containing vacuole (SCV) (Cirillo et al., 1998; Feng et al., 2003; Kuhle & Hensel, 2004; A. K. Lee et al., 2000; Ochman et al., 1996; Shea et al., 1996).

Regulation of the SPI-2 pathogenicity island is complex and involves silencing by the nucleoid-associated protein H-NS (Gao et al., 2015, 2017; Liu et al., 2010; Lucchini et al., 2006; Winardhi et al., 2015) and anti-silencing by response regulators (Desai et al., 2016; Walthers et al., 2011; Will et al., 2014). Response regulators are part of a signal transduction system prevalent in bacteria. Such two-component systems consist of a membrane-bound histidine kinase and a cytoplasmic response regulator (RR), which binds to DNA and activates gene transcription (Hoch & Silhavy, 1995). The SsrA/B system plays a crucial role in regulating SPI-2 gene expression (Feng et al., 2003; Gao et al., 2017; Kenney, 2019; Lee et al., 2000). Activation of SPI-2 genes requires phosphorylation of the SsrB RR on a conserved aspartic acid residue by its kinase SsrA (Carroll et al., 2009; Feng et al., 2004). Upon activation, SsrB binds to AT-rich regions of DNA and activates transcription of SPI-2 promoters via displacement of the nucleoid-binding protein H-NS (Walthers et al., 2011), as well as direct recruitment of RNA polymerase (Walthers et al., 2007). The expression of *ssrAB* is surprisingly complex; a promoter for *ssrB* resides in the coding region of *ssrA*, a 30 bp intergenic region lies between *ssrA* and *ssrB*, and both genes have extensive untranslated regions, suggesting post-transcriptional or translational control (Feng et al., 2003). Each component of the enigmatic SsrA/B system is regulated by separate global regulators EnvZ/OmpR (Feng et al., 2003; A. K. Lee et al., 2000) and PhoQ/P (Bijlsma & Groisman, 2005), indicating an uncoupling of the operon. This complexity was confounding, but recent studies demonstrated a non-canonical role for unphosphorylated SsrB in the absence of its kinase SsrA in driving biofilm formation and establishment of the carrier state, indicating a dual function for SsrB in controlling *Salmonella* lifestyles (Desai et al., 2016; Desai & Kenney, 2017). Recently, we counted SsrA and SsrB molecules using photoactivation localization microscopy (PALM) and demonstrated their uncoupling and stimulation by acid pH (Liew et al., 2019). This complex hierarchy of gene activation ensures that activation of SPI-2 occurs only under conditions that presumably mimic the macrophage vacuole such as low pH, low Mg^2+^ and high osmolality (Chakraborty et al., 2015; Choi & Groisman, 2016; Deiwick et al., 1999; Miao et al., 2002).

Upon encountering the acidic environment of the vacuole, *Salmonella* acidifies its cytoplasm in an OmpR-dependent manner through repression of the *cadC/BA* system (Chakraborty et al., 2015). Intracellular acidification provides an important signal for expression and secretion of SPI-2 effectors. There is now increasing evidence that this change in intracellular pH is important for pathogenesis (Chakraborty et al., 2015, 2017; Choi & Groisman, 2016; Kenney, 2019; Liew et al., 2019), although little is known as to how cytoplasmic acidification leads to SPI-2 gene activation. In particular, the effect of acidification on SsrB has not been thoroughly investigated until now.

In this study, we demonstrate that acid-stimulated DNA binding by SsrB is not a property of the C-terminal DNA binding domain, but is driven by a single conserved histidine residue in the receiver domain. A mutant substituting histidine 12 (His12) with a glutamine residue retained 100% of the wild-type transcriptional activity at neutral pH and eliminated activity at acid pH. Substitution of His12 with an aromatic amino acid retained pH sensitivity, but substantially reduced its activity compared to the wild-type. In addition to influencing SsrB pH sensing, His12 also contributed to SsrB phosphorylation. Eliminating acid sensitivity renders *Salmonella* avirulent and thus represents a potential target for controlling infection.

## RESULTS

### Acid pH increases SsrB affinity for DNA-

Previous studies used single particle tracking PALM (spt-PALM) and demonstrated an acid-dependent increase in SsrB binding to DNA in single cells (Liew et al., 2019). Hence, we were interested in determining the precise changes of a SPI-2 promoter containing a known SsrB binding site in response to acid pH. The SPI-2 promoter *sseI* (formerly known as *srfH*) contains an SsrB binding site of ∼ 47 bp, as determined by DNase I footprinting (Feng et al., 2004). We used this 47 bp region to construct a DNA hairpin in order to measure SsrB binding affinity using a single molecule unzipping assay ((Gulvady et al., 2018), See Methods). We compared SsrB binding at neutral pH (7.4) and acid pH (6.1), the pH that we determined was the intracellular pH (pH_i_) of *Salmonella* in the SCV (Chakraborty et al., 2015, 2017). At pH 7.4, the binding of SsrB to *sseI* was extremely cooperative (Hill coefficient (*n*) = 7.64 ± 1.2) and the dissociation constant (*K*_*D*_) was ∼147.8 ± 1.96 nM (**Fig. 1a**). At acid pH, the Hill coefficient was relatively unchanged (*n* = 11.7 ± 5.5), while the *K*_*D*_ decreased substantially, to 4.64 ± 0.19 nM (**Fig. 1b**). Hence, at the SPI-2 promoter *sseI*, SsrB binding was ∼32 times higher affinity at acid pH than at neutral pH and binding at both neutral and acid pH was highly cooperative.

**Fig. 1:**
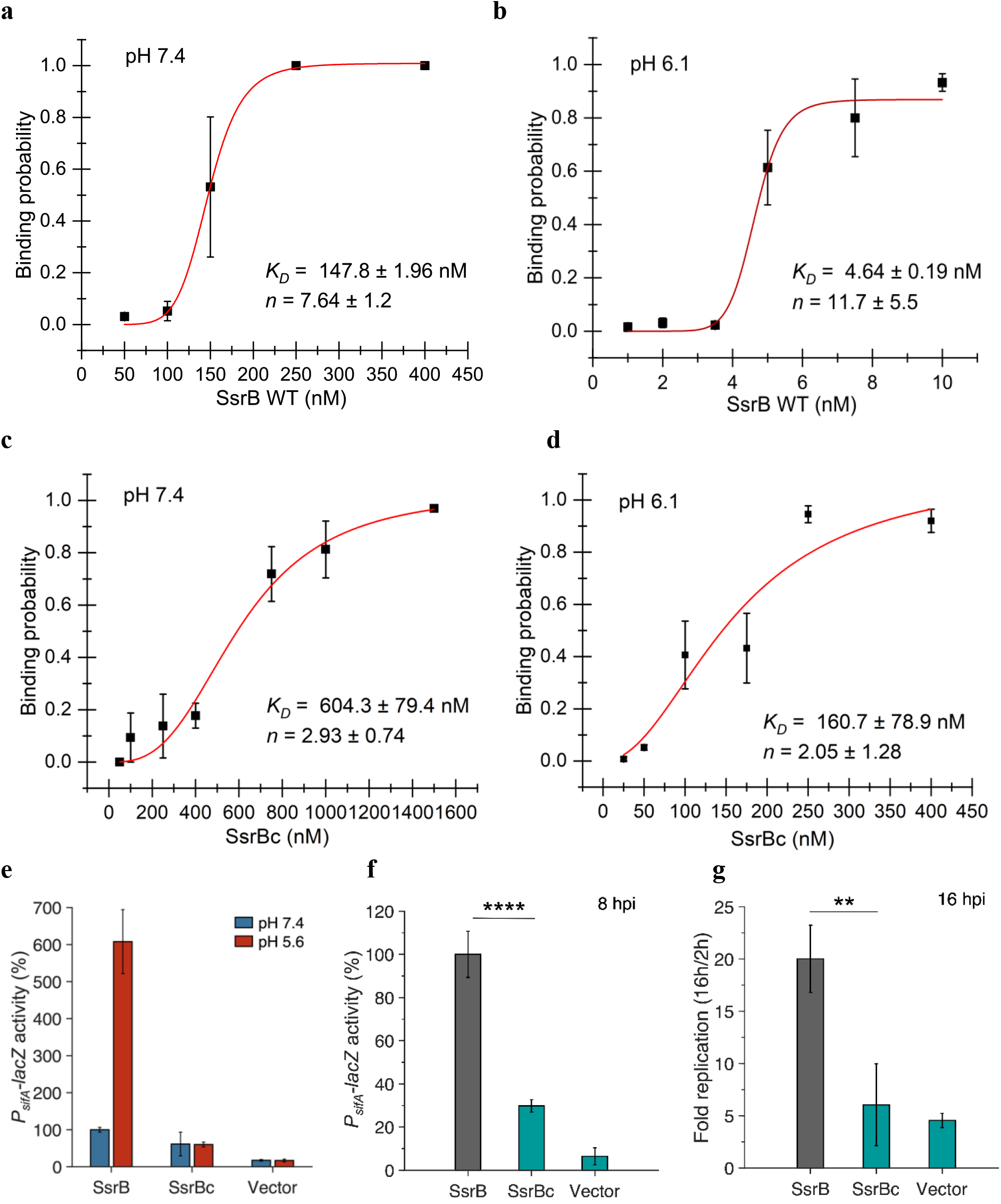
SsrBc is not the locus of acid-sensitive DNA binding. **a**, The plots represent the binding probability of SsrB or SsrBc to the *sseI* DNA hairpin as a function of protein concentration (nM) at pH 7.4 and 6.1. At neutral pH, SsrB binds to the *sseI* promoter with a *K*_*D*_ of 147.8 ± 1.96 nM, Hill coefficient (*n*)= 7.64 ± 1.2. **b**, At acid pH 6.1, the *K*_*D*_ was 4.64 ± 0.19 nM and *n* = 11.7 ± 5.5. (c) At neutral pH, SsrBc binds to the *sseI* promoter with a *K*_*D*_ of 604.3 ± 79.4 nM, Hill coefficient (*n*) = 2.93 ± 0.74. **d**, At acidic pH 6.1, the *K*_*D*_ was 160.7 ± 78.9 nM and *n* = 2.05 ± 1.28. The error bars represent the standard deviation of three to five independent measurements. The red line represents the curve derived from fitting the points to the Hill equation to determine the *K*_*D*_. The absence of error bars indicates that the standard deviation was < the symbol. **e**, *P*_*sifA*_*-lacZ* activity in the presence of SsrB or SsrBc grown in MGM media pH 7.4 (blue bars) and pH 5.6 (red bars) after 3 hours of 0.1% arabinose induction. In comparison to SsrB activity at pH 7.4, the activity of the SsrBc strains were reduced to 61% and 60% activity in pH 7.4 and 5.6 media, respectively. **f**, *P*_*sifA*_*-lacZ* activity measured from strains recovered at 8 h post HeLa infection. The activity for the SsrBc strain was 30% of the wildtype SsrB strain (****=p<0.0001, n=3) **g**, During HeLa cell infection, the wildtype full-length SsrB strain increased 20-fold after 16 hpi. The SsrBc strain only increased 6-fold over the same time period (**=p<0.01, n=3).

### Acid-stimulated DNA binding does not reside in the DNA binding domain of SsrB (SsrBc)-

SsrB has an N-terminal receiver domain that is phosphorylated and a C-terminal DNA binding domain, connected by a flexible linker. SsrBc, comprising the linker and DNA binding domain of SsrB (residues 138-212), has been shown to induce the expression of SPI-2 genes *in vitro* at a level similar to the full-length SsrB (Feng et al., 2004). To determine if SsrBc alone was responsible for the pH-dependent DNA binding, we used the single molecule unzipping assay to measure SsrBc binding to the 47 bp *sseI* DNA hairpin at neutral and acid pH. At pH 7.4, the *K*_*D*_ of SsrBc was 604.3 ± 79.4 nM, with a Hill coefficient of 2.93 ± 0.74 (**Fig. 1c**). Thus, the isolated C-terminus of SsrB binds to DNA with a 4-fold lower affinity and a reduction in cooperativity compared to the full-length protein. At pH 6.1, the *K*_*D*_ value decreased to 160.7 ± 78.9 nM, without a substantial change in cooperativity (*n* = 2.05 ± 1.28) (**Fig. 1d**). Hence, in comparison to full-length SsrB, SsrBc binds DNA with reduced affinity and cooperativity at both neutral and acid pH. Although the affinity of SsrBc for DNA increased at acid pH, the decrease in *K*_*D*_ was only 4-fold, whereas the full-length SsrB demonstrated a 32-fold decrease over this same pH range. The change in DNA binding at acid pH with SsrBc was similar to the fold change of full-length SsrB at the non-SPI-2 promoter *csgD* (Liew et al., 2019).

SsrBc was also incapable of supporting acid-stimulated SPI-2 transcriptional activity compared to full-length SsrB both *in vitro* and *in vivo*. Transcriptional activity of the *sifA* promoter (another SPI-2-regulated promoter) was measured in a Δ*ssrB* strain (14028s *ΔssrB attB::pAH125 P*_*sifA*_*-lacZ*, DW637) carrying SsrB or SsrBc on a plasmid under the regulation of an inducible arabinose promoter. Early log phase cells grown in SPI-2 non-inducing (pH 7.4) and inducing conditions (pH 5.6) were induced with 0.1% arabinose, and the *β*-galactosidase activity was measured after 3 hours. All activities henceforth described are expressed as the relative P_*sifA*_*-lacZ* activity of a given SsrB mutant with respect to P_*sifA*_*-lacZ* activity of the SsrB wild-type expressing strain grown in MGM pH 7.4. For cells grown at pH 7.4, the relative activity of *P*_*sifA*_*-lacZ* in the presence of SsrBc or SsrB was 61% and 100%, respectively. In cells grown at pH 5.6, the activities were 60% and 570%, respectively, relative to the wild-type SsrB activity at pH 7.4 (**Fig. 1e**). Thus, while SsrB exhibited a ∼5.7-fold increase in transcriptional activity when grown in acid pH, transcription by SsrBc was not stimulated at acid pH (0.92-fold). Inside HeLa cells, the SsrBc-expressing strain exhibited reduced intracellular survival (16 hpi) and decreased *P*_*sifA*_*-lacZ* activity (8 hpi) compared to SsrB. Intracellular survival of the SsrBc strain was 30% and *P*_*sifA*_*-lacZ* activity was 30% compared to the wildtype SsrB strain (**Fig. 1f,g, Supplementary Fig. 3**). Thus, the isolated C-terminal domain was insufficient to support intracellular survival and replication *in vivo*.

### Analysis and comparison of the NarL/FixJ sub-family of response regulators-

Of all of the NarL/FixJ sub-family of response regulators (RRs), SsrB is the only member for which pH-dependence of DNA binding has been reported (Liew et al., 2019). The RRs RcsB and EvgA are involved in acid stress responses in *E. coli*, while other RRs have functions that do not directly rely on changes in pH (Belcheva & Golemi-Kotra, 2008; Boon & Dick, 2002; Castanié-Cornet et al., 2010; Jordan et al., 2006; Nishino et al., 2003; Olmedo et al., 1990; Stewart, 1994). A comparison of NarL/FixJ RRs reveals that SsrB has the highest isoelectric point in this group (*pI* = 7.12), whereas most of the others range from 4.95 to 6.85 (**Table 1**). Therefore, an increase in the overall protonation of SsrB as the bacterial cytoplasmic pH decreases within the SCV could contribute to the increase in DNA binding affinity that we observed (**Fig. 1b**). Because SsrBc has an even higher *pI* of ∼9.36, it is unlikely that it would undergo enough change in protonation to confer pH sensitivity under SPI-2 inducing conditions, as we also observed (**Fig. 1c-e**).

**Table 1.** Comparison of NarL/FixJ sub-family of response regulators

| Protein | Organism | pI | Number of Histidines | %Identity with SsrB |
| --- | --- | --- | --- | --- |
| SsrB | *Salmonella* Typhimurium | 7.12 | 9 | 100.0 |
| RcsB | *E. coli* K12 | 6.85 | 3 | 25.47 |
| RcsB | *Salmonella* Typhimurium | 6.85 | 3 | 25.47 |
| EvgA | *Shigella flexneri* | 6.83 | 3 | 28.16 |
| Spo0A | *Bacillus stearothermophilus* | 6.31 | 9 | 25.24 |
| NarL | *E. coli* K12 | 5.73 | 6 | 28.30 |
| DegU | *Bacillus subtilis* | 5.66 | 11 | 26.75 |
| DosR | *Mycobacterium tuberculosis* | 5.62 | 1 | 27.54 |
| VraR | *S. aureus* | 5.47 | 5 | 29.25 |
| StyR | *Pseudomonas fluorescens* | 5.42 | 6 | 26.47 |
| LiaR | *Enterococcus faecium* | 5.11 | 5 | 29.95 |
| Spr1814 | *Streptococcus pneumoniae* | 4.95 | 1 | 25.60 |
| PhoP | *E. coli* | 5.10 | 6 | 22.49 |
| OmpR | *E. coli* | 6.04 | 3 | 31.62 |

In addition to its high *pI*, SsrB also contains a greater number of histidine residues (9) compared to other RRs. Histidine residues are known to play a role in pH sensitivity of numerous proteins (Furman et al., 2015; Mulder et al., 2015; Müller et al., 2009; Tu et al., 2009), as the *pK*_*a*_ of the histidine side chain (∼6.45) enables it to protonate under the physiological pH range (Platzer et al., 2014). The pH threshold of the *Salmonella* cytoplasm below which it expressed and secreted SPI-2 effectors was determined to be between 6.45 and 6.7 (Chakraborty et al., 2017; Kenney, 2019). Hence, we reasoned that the high number of histidines in SsrB potentially contributed to its acid-stimulated DNA binding. Since SsrBc alone does not show a very strong pH-dependence (**Fig. 1c-e**), we screened histidine residues in the N-terminal phosphorylation domain for candidates likely to contribute to the pH sensitivity. We identified four histidine residues in the receiver domain, three of which were unique to SsrB (His28, His34 and His72, **Supplementary Fig. 1**). The fourth histidine residue, His12, was well-conserved amongst most of the NarL/FixJ RRs (**Supplementary Fig. 1)**. Since none of these other RRs had been shown to be pH sensitive, we initially screened the three unique histidine residues for pH sensitivity.

### Unique histidines in the receiver domain do not affect pH sensitivity

In an i-TASSER predicted structure of the receiver domain of SsrB (Roy et al., 2010; Yang et al., 2015; Zhang, 2008), the three unique histidine residues appear to be solvent exposed, with His28 present in the loop between *α1* and *β2*, His34 on *β3* and His72 on *α3* (**Supplementary Fig. 2**). To assess the effect of these N-terminal histidine residues on SsrB pH sensitivity, each histidine was substituted with alanine using PCR mutagenesis. The single histidine SsrB mutants SsrB H28A, SsrB H34A and SsrB H72A were assayed for their SPI-2 transcriptional activity when grown in MGM at pH 7.4 or 5.6. There were no substantial differences in the acid-stimulated increase in *P*_*sifA*_-*lacZ* activity of any of the singly substituted mutant strains compared to wild-type SsrB (**Fig. 2a,b**). The increase in *P*_*sifA*_-*lacZ* activity at acid pH vs neutral pH for SsrB H28A, SsrB H34A and SsrB H72A was 4.2, 4.8 and 4.5-fold respectively, similar to the increase of the wild-type (5-fold).

**Fig. 2:**
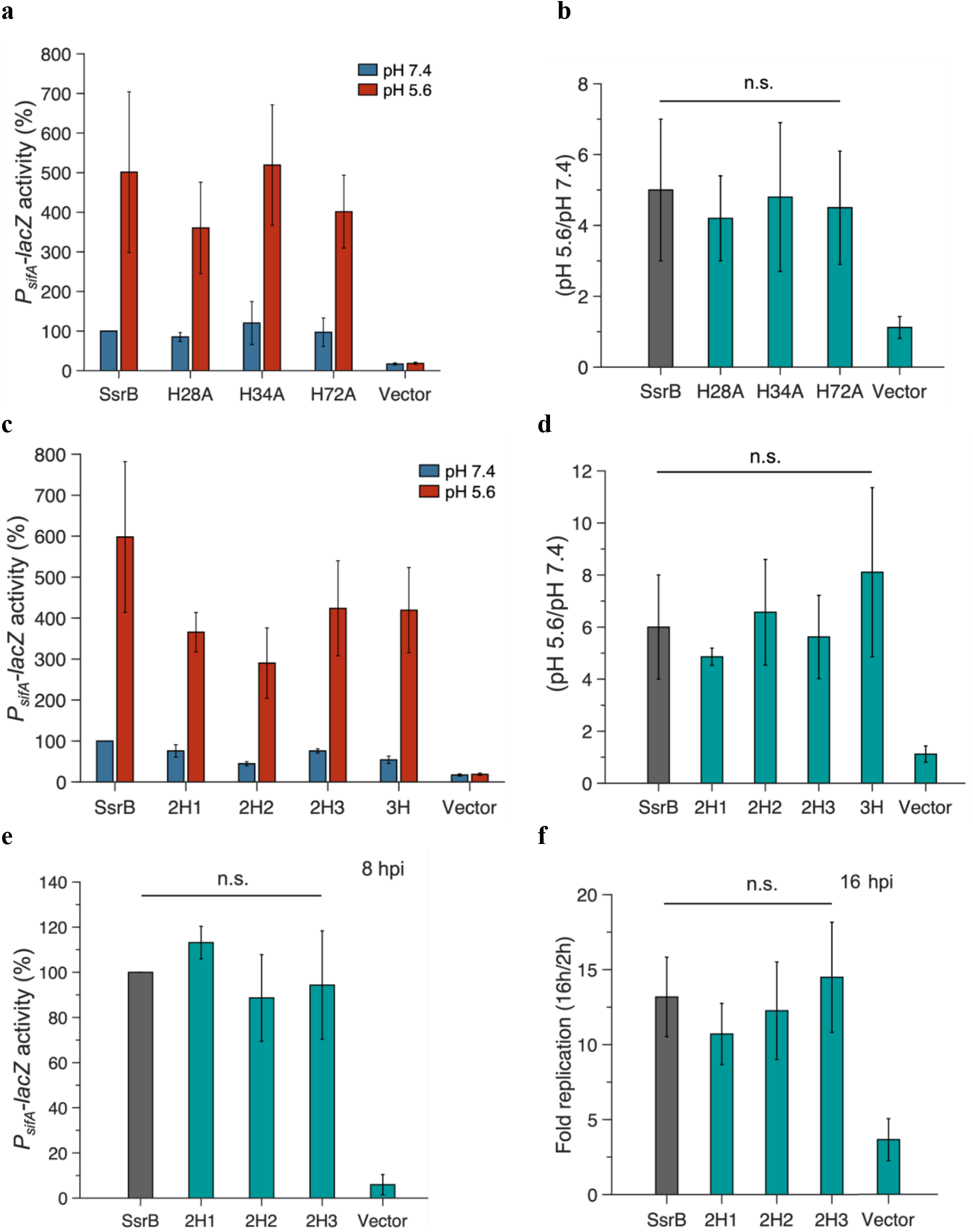
Non-conserved histidines in the receiver domain do not confer acid-sensitivity to SsrB. **a**, *P*_*sifA*_*-lacZ* activity in the presence of SsrB, single histidine mutants of SsrB or vector only control grown in MGM media pH 7.4 (blue bars) and pH 5.6 (red bars) after 3 hours of 0.1% arabinose induction. **b**, The increase in activity at pH 5.6 shown by SsrB H28A, SsrB H34A and SsrB H72A is 4.2, 4.8 and 4.5-fold respectively, which is comparable to the wild-type SsrB (5-fold) (n.s.=p>0.05, n=3). **c**, *P*_*sifA*_*-lacZ* activity in the presence of SsrB or double or triple histidine mutants of SsrB grown in MGM media pH 7.4 (blue bars) and pH 5.6 (red bars) after 3 hours of 0.1% arabinose induction. **d**, The increase in activity at pH 5.6 shown by SsrB 2H1, SsrB 2H2 and SsrB 2H3 (4.8, 6.6, 5.6 and 7.3-fold respectively), was comparable to the increase in activity shown by wild-type SsrB (5.9-fold) (n.s.=p>0.05, n=3). **e**, *P*_*sifA*_*-lacZ* activity measured from strains recovered at 8 h post HeLa infection. The activity of the SsrB 2H1, SsrB 2H2 and SsrB 2H3 expressing strains was 113.1±7.2%, 88.7±19.2% and 94.4±24%, respectively, relative to the SsrB expressing strain (n.s.=p>0.05, n=3). **f**, Intracellular survival of strains in HeLa cells at 16 hpi. During HeLa cell infection, the c.f.u./ml of the SsrB expressing strain increased 13.2-fold at 16 hpi. For SsrB 2H1, SsrB 2H2 and SsrB 2H3 expressing strains, the c.f.u./ml increased 12.7, 13.3 and 14.5-fold, respectively, similar to SsrB (n.s.=p>0.05, n=3).

The double histidine mutants SsrB H28A-H34A (SsrB 2H1), SsrB H28A-H72A (SsrB 2H2) and SsrB H34A-H72A (SsrB 2H3) and triple histidine mutant SsrB H28A-H34A-H72A (SsrB 3H) similarly showed no substantial differences in acid-stimulation of *P*_*sifA*_-*lacZ* activity *in vitro*. The increase in *P*_*sifA*_-*lacZ* activity at acid vs neutral pH for strains expressing SsrB 2H1, SsrB 2H2, SsrB 2H3 and SsrB 3H (4.8, 6.6, 5.6 and 7.3-fold, respectively) was similar to the increase by the wild-type (5.9-fold) (**Fig. 2c,d**). *In vivo*, when the strains expressing SsrB 2H1, SsrB 2H2 or SsrB 2H3 were used to infect HeLa cells, the *P*_*sifA*_-*lacZ* activity at 8 hpi and subsequent survival at 16 hpi was also comparable to the wild-type SsrB. The *P*_*sifA*_-*lacZ* activity measured for SsrB 2H1, 2H2 and 2H3 at 8 hpi was 113%, 89%, 94% relative to the wild-type (**Fig. 2e**). Subsequently, intracellular survival at 16 hpi was comparable to the wild-type survival (13.2-fold) (**Fig. 2f**).

### Highly conserved Histidine 12 is essential for acid-stimulated DNA binding of SsrB-

As substitutions of unique histidines failed to abolish the pH sensitivity of SsrB, we screened the remaining N-terminal histidine residue, His12, for pH sensitivity. His12 lies in the loop between β1 and α1 in the receiver domain (**Fig. 3a**). It is in close proximity to the phosphorylated residue, Asp56, the metal coordinating residues Asp10 and Asp11, and the polar contact residue Lys106 (**Fig. 3a**). We compared the *P*_*sifA*_-*lacZ* activity of alanine and asparagine substitutions at position 12 to wild-type SsrB, they were only 66% and 77% active at pH 7.4 (**Fig. 3b)**. At pH 5.6, their activity increased by only 1.8- and 1.5-fold, respectively (**Fig. 3b,c**). A glutamine substitution at position 12 was 98.9% active at pH 7.4, while at pH 5.6, there was no increase in activity (0.93-fold). These results indicate that the conserved His12 is the major driver of the acid-stimulated DNA binding of SsrB.

**Fig. 3:**
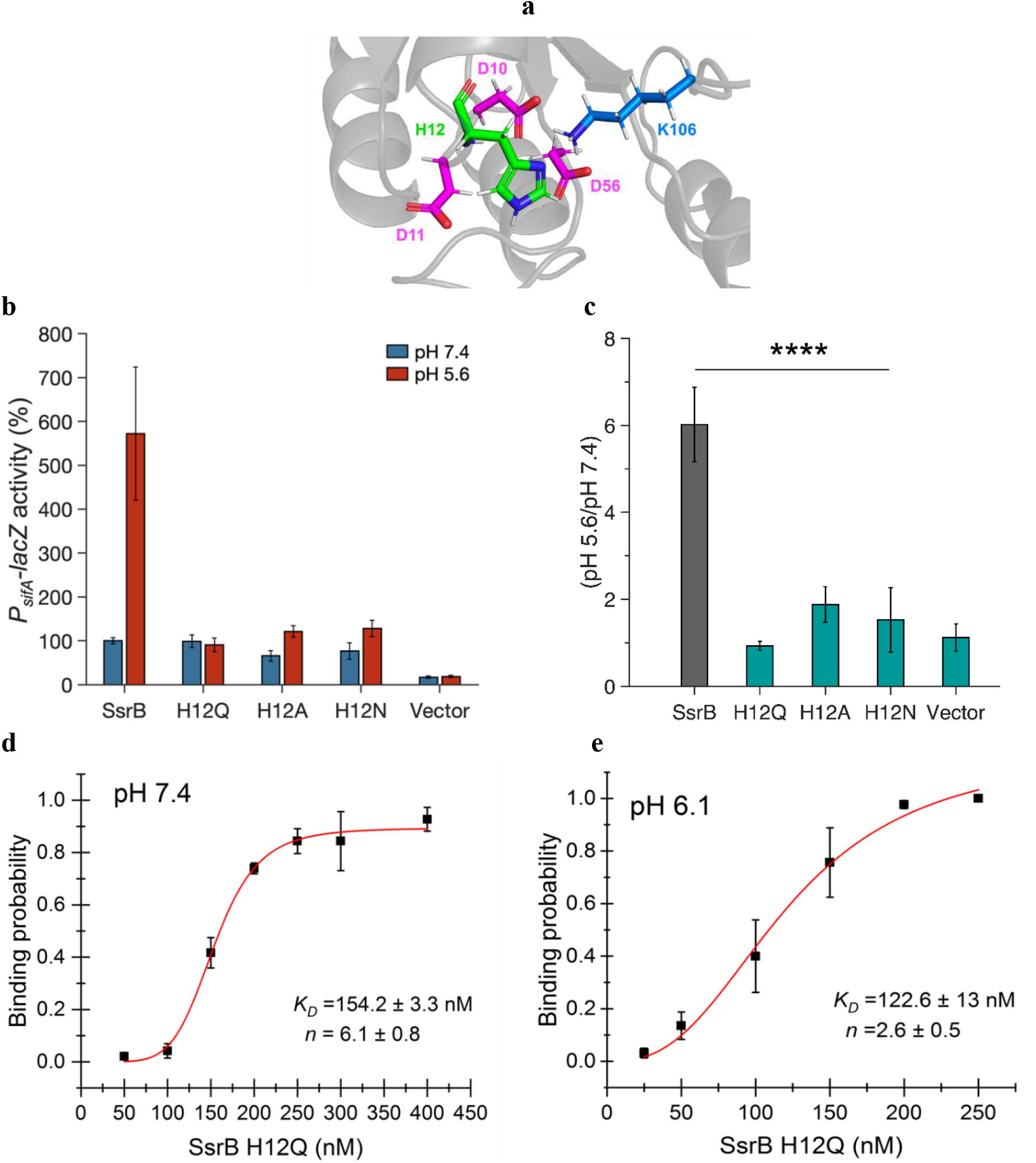
Histidine 12 is essential for acid-stimulated DNA binding of SsrB. **a**, The location of His12, on a predicted receiver domain structure of SsrB (visualized with PyMol). His12 is in the vicinity of the active site residues Asp10, Asp11, Asp56 and Lys106. **b**, *P*_*sifA*_*-lacZ* activity in the presence of SsrB or H12 mutants of SsrB grown in MGM media pH 7.4 (blue bars) and pH 5.6 (red bars) after 3 hours of 0.1% arabinose induction. **c**, The H12A, H12Q and H12N substitutions showed no increase in activity at pH 5.6 (1.8, 0.93, and 1.5-fold respectively), compared to wild-type SsrB (6-fold) (p<0.0001, n=3). **d**, and **e**, The plots represent the binding probability of SsrB H12Q to the *sseI* DNA hairpin as a function of SsrB H12Q concentration (nM) at pH 7.4 and 6.1. **d**, At neutral pH, SsrB H12Q binds to the *sseI* promoter with a *K*_*D*_ of 154.2 ± 3.3 nM, *n* = 6.1 ± 0.8. **e**, At acidic pH 6.1, the *K*_*D*_ was 122.6 ± 13 nM and *n* = 2.6 ± 0.5. The error bars represent the standard deviation of three to five independent measurements. The red line represents the curve derived from fitting the points to the Hill equation to determine the *K*_*D*_. The absence of error bars indicates that the standard deviation was < the symbol (n=3).

As the glutamine substitution retained full activity at pH 7.4, we used purified SsrB H12Q to measure the effect of His12 substitution on the DNA binding activity of SsrB in the single molecule unzipping assay. At pH 7.4, the *K*_*D*_ of binding of SsrB H12Q to the *sseI* promoter was 154.2 ± 3.3 nM with a Hill coefficient of 6.1 ± 0.8 (**Fig. 3d**), similar to wild-type SsrB. At pH 6.1, the *K*_*D*_ remained relatively unchanged at 122.6 ± 13 nM, while the cooperativity was reduced to 2.6 ± 0.5 (**Fig. 3e**). This value was similar to the cooperativity of SsrBc (**Fig. 1c,d**). Hence, a substitution at His12 eliminated both the increase in DNA binding affinity and the large change in cooperativity of SsrB at acid pH.

### A role for the aromatic ring in pH sensitivity-

To elucidate the mechanism by which His12 contributes to pH sensitivity, additional amino acids were substituted in place of histidine. Histidine can act both as a basic and an aromatic amino acid, hence it can have a variety of interactions, depending on its protonation state (Liao et al., 2013). The protonated form of histidine at low pH would theoretically be mimicked by substitution with a positively charged amino acid, while substitution with an aromatic amino acid could mimic the uncharged aromatic imidazole ring. We therefore made lysine, tyrosine and phenylalanine substitutions at His12 and screened the activity and pH sensitivity of these mutants. Both aromatic substitutions exhibited substantial reductions in their activity but maintained pH-sensitivity. SsrB H12K *P*_*sifA*_-*lacZ* activity was 33.5% of wild-type SsrB at pH 7.4 (**Fig. 4a**) and its activity only increased 2.7-fold at acid pH (**Fig. 4b**). Activity of H12Y and H12F strains was reduced to 29.5% and 24.2%, respectively (**Fig. 4a**), but at pH 5.6, these mutants showed a 6.9-fold and 6.7-fold increase in activity, which was comparable to the wild-type (**Fig. 4b**). Hence the lysine substitution at His12 reduced both activity and pH sensitivity in SsrB, while the tyrosine and phenylalanine substitutions led to a reduction in the activity but retained the acid stimulation, indicating a role for the imidazole aromatic ring in the acid stimulation of transcriptional activity.

**Fig. 4:**
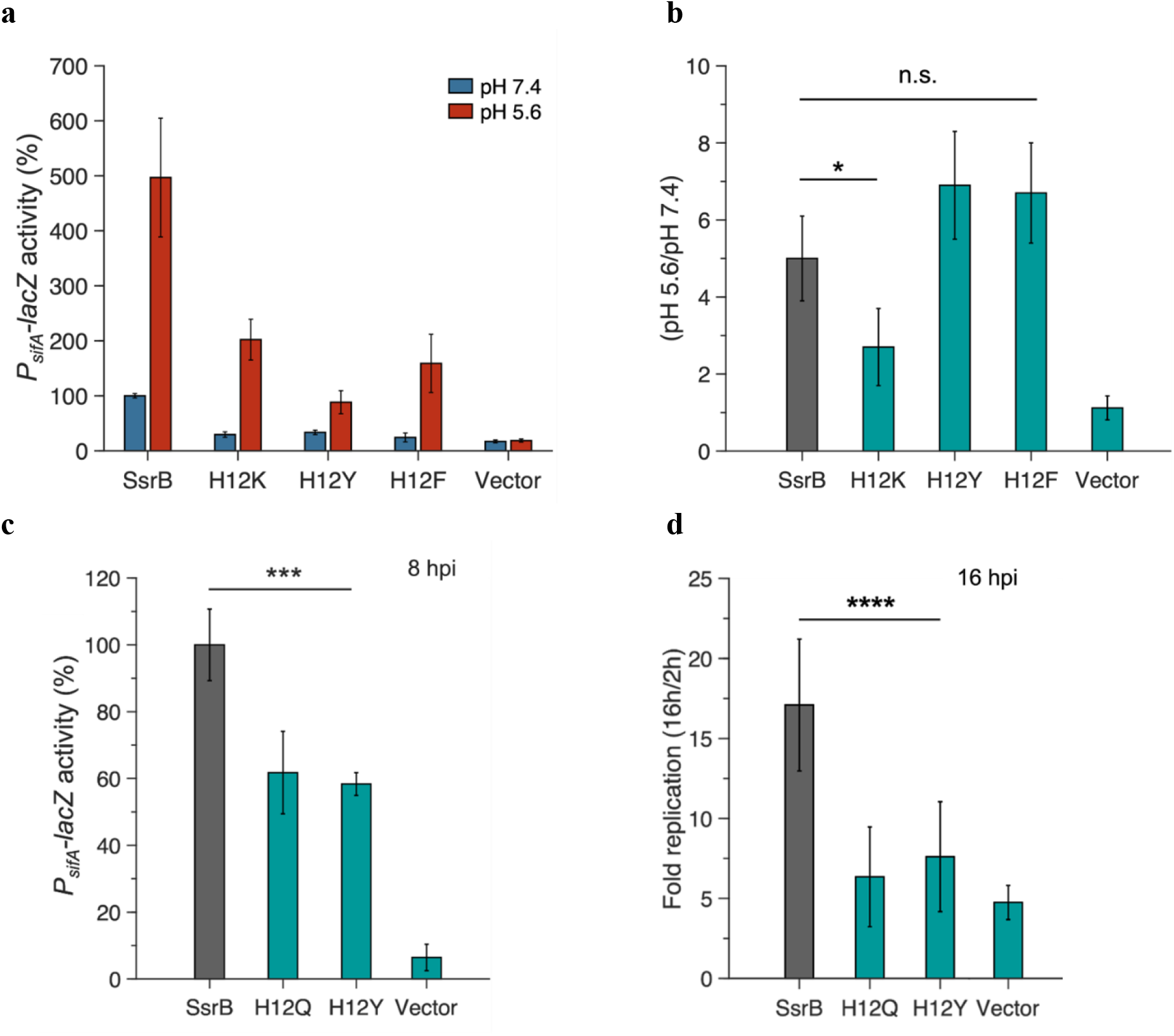
Aromatic substitutions at position 12 retain SsrB pH sensitivity. **a**, *P*_*sifA-lacZ*_ activity in the presence of SsrB or H12 mutants of SsrB grown in MGM media pH 7.4 (blue bars) and pH 5.6 (red bars) after 3 hours of 0.1% arabinose induction. The H12K, H12Y and H12F substitutions showed a decrease in activity at pH 7.4 compared to SsrB (33.5%, 29.5% and 29.2%). **b**, SsrB H12K only showed a moderate increase in activity at pH 5.6 (2.7-fold), however the SsrB H12Y and H12F substitutions retained acid-stimulated increase in the activity (6.9 and 6.7-fold respectively), comparable to the wild-type (5-fold). **c**, *P*_*sifA*_*-lacZ* activity measured from strains recovered at 8 h post HeLa infection. The activity of the SsrB H12Q and SsrB H12Y expressing strains was 61.7±12% and 58.3±3.4%, respectively, relative to the SsrB expressing strain (p<0.001, n=3). **d**, Intracellular survival of strains in HeLa cells at 16 hpi. During HeLa cell infection, the c.f.u./ml of the SsrB expressing strain increased 19-fold at 16 hpi. For SsrB H12Q and SsrB H12Y expressing strains, the c.f.u./ml only increased 6.35 and 7.6-fold, respectively (p<0.0001, n=3).

Substitutions at His12 also led to a reduction in intracellular *P*_*sifA*_*-lacZ* activity and survival of *Salmonella* infections of HeLa cells. At 8 hpi, the *P*_*sifA*_*-lacZ* activity of SsrB H12Q and H12Y expressing strains was reduced to 61% and 58% compared to SsrB (**Fig. 4c**). Similarly, at 16 hpi, intracellular survival of H12Q and H12Y was reduced to 33.4% and 40% of the wild-type (**Fig. 4d, Supplementary Fig. 3**). Although the H12Y substitution retained acid pH-sensitivity *in vitro*, the low intracellular transcriptional activity and survival can be attributed to the overall reduction in its activity compared to the wild type.

### Substitution of His12 reduces SsrB phosphorylation and eliminates pH sensitivity-

As SsrB phosphorylation is essential for SPI-2 activation (Feng et al., 2004) and His12 is proximal to the phosphorylated residue Asp56 (**Fig. 3a**), we examined the effect of His12 substitution on SsrB phosphorylation. SsrB is readily phosphorylated *in vitro* by the small molecule phosphodonor phosphoramidate (PA) (Feng et al., 2004), we thus compared the phosphorylation of wild-type SsrB with the H12Q mutant. The amount of phosphorylated protein (SsrB∼P or H12Q∼P) generated was determined by resolving the reaction mixture on a C_4_ reverse phase HPLC column under a 40-50% acetonitrile gradient (Feng et al., 2004).

12 μM of SsrB or H12Q was phosphorylated with 2.5 mM PA at pH 7.4 for varying amounts of time. About 89.8% SsrB was phosphorylated at saturation (*B*_*max*_), with a *K*_*obs*_ of 0.17 min^-1^ (**Fig. 5a**). By comparison, only 44.8% H12Q was phosphorylated at saturation, with a reduced *K*_*obs*_ of 0.096 min^-1^ (**Fig. 5b**). As phosphorylation of SsrB was much faster than H12Q, a new reaction varied the amount of PA during a 10 min incubation. For wild-type SsrB, ∼2.9 mM PA was sufficient to phosphorylate 50% of the protein (*K*_*M*_) at pH 7.4 (**Fig. 5c**). At pH 6.1, the amount of PA required to phosphorylate 50% of the protein increased to 13.4 mM (**Fig. 5d**). Hence the efficiency of SsrB phosphorylation was reduced 4.6-fold at acid pH (**Fig. 5g**). SsrB H12Q required a higher concentration of PA (*K*_*M*_ = 7.1 mM) to phosphorylate 50% of the protein at pH 7.4 (**Fig. 5e**), but at pH 6.1, the *K*_*M*_ was relatively unchanged at 5.05 mM PA (**Fig. 5f**). Hence, substituting His12 both reduced SsrB phosphorylation at neutral pH and abolished pH-dependent changes in SsrB phosphorylation (**Fig. 5g**). An interesting observation was the delayed retention time of H12Q as compared to SsrB, irrespective of the pH of the phosphorylation buffer, suggestive of conformational differences between the two proteins (**Supplementary Fig. 6**).

**Fig. 5:**
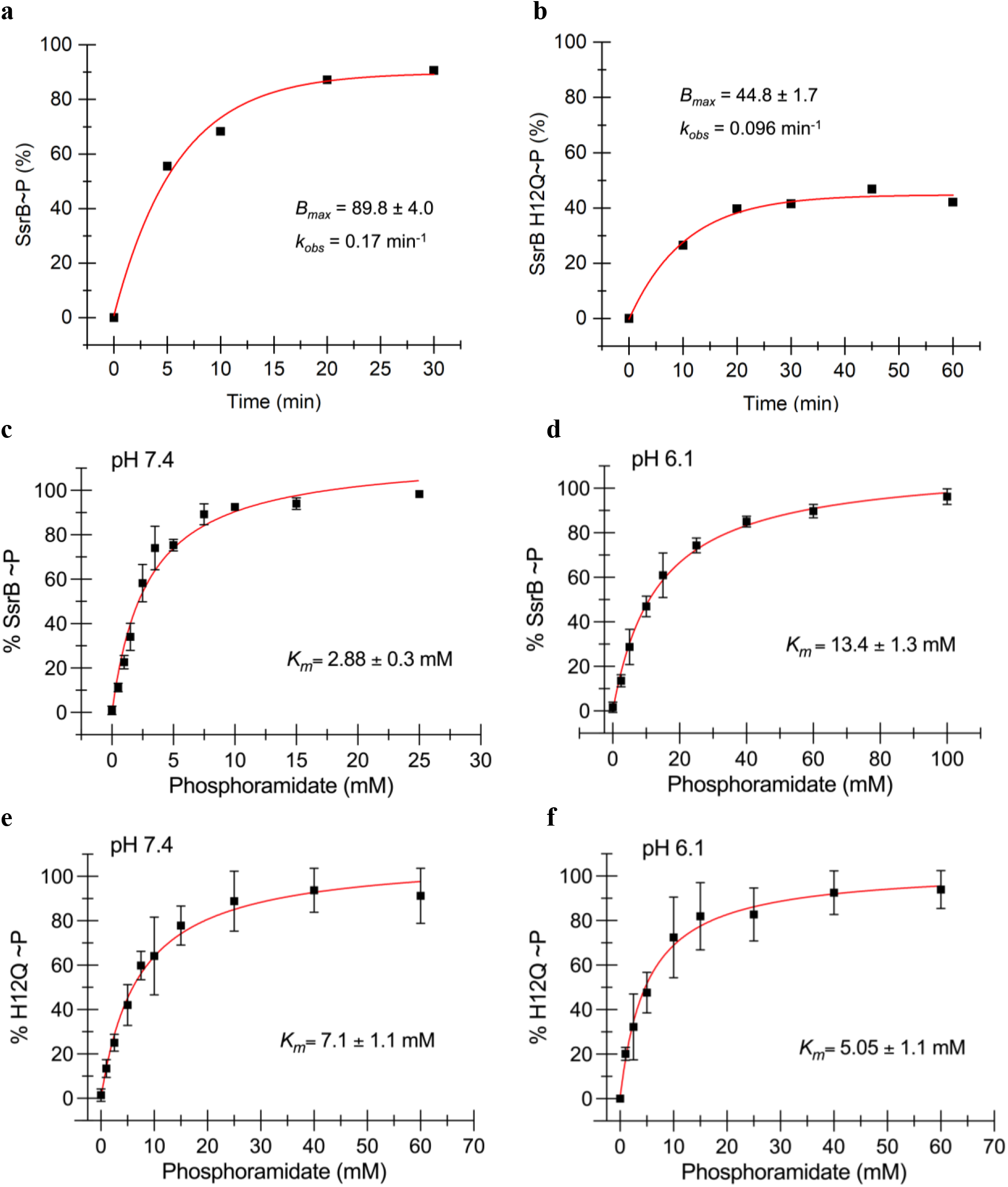
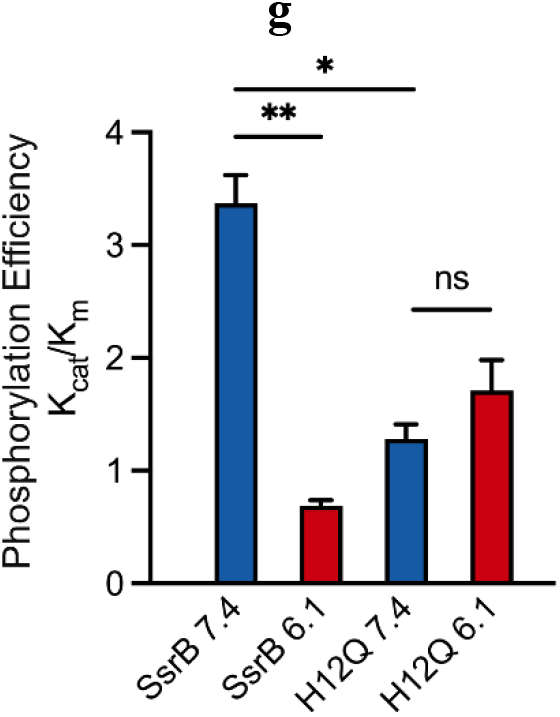
His12 substitution affects SsrB phosphorylation. The plots represent the percentage of phosphorylated protein against the reaction time (**a** and **b**) or the amount of phosphoramidate (PA) in the phosphorylation reaction (**c**-**f**). **a**, A phosphorylation reaction with 12 μM SsrB and 2.5 mM PA yielded 89.8% ± 4.0 phosphorylated protein at saturation (*B*_*max*_) with a *K*_*obs*_ of 0.17 min^-1^. **b**, A phosphorylation reaction with 12 μM H12Q and 2.5 mM PA yielded 44.8 ± 1.7% phosphorylated protein at saturation with a *K*_*obs*_ of 0.096 min^-1^. The red line represents the curve derived from fitting the points to a single exponential decay model. **c**, The amount of PA required to phosphorylate 50% of the protein, *K*_*M*_, for SsrB phosphorylation was 2.88 ± 0.3 mM PA at pH 7.4 and **d**, 13.4 ± mM at pH 6.1. **e**, The *K*_*M*_ for SsrB H12Q phosphorylation was 7.1 ± 1.1 mM at pH 7.4, and **f**, 5.05 ± 1.1 mM at pH 6.1. The error bars represent the standard deviation of two independent measurements. The red line represents the curve derived from fitting the points to the hyperbolic equation to determine the *K*_*M*_. The absence of error bars indicates that the standard deviation was < the symbol. **g**, A comparison of the phosphorylation efficiencies (*K*_*cat*_ / *K*_*M*_) of SsrB and H12Q at pH 7.4 and 6.1. The efficiency of SsrB phosphorylation was reduced by 4.6-fold at pH 6.1. At pH 7.4, H12Q phosphorylation efficiency reduced 2.5-fold compared to SsrB. There was no significant difference between the phosphorylation efficiency of H12Q at 7.4 and pH 6.1 (** = p<0.01, * = p<0.05, n.s.= p>0.05, n=2).

## DISCUSSION

While the effector domains of response regulators confer target promoter specificity, it is the receiver domains that fine-tune their activation in response to environmental signals. In the case of SsrB, the receiver domain senses a reduction in pH to allosterically increase DNA binding affinity of the C-terminal effector domain. SsrBc alone had a 4-fold lower DNA binding affinity than SsrB, and this affinity only increased by 4-fold in acidic pH *in vitro* (**Fig. 1c,d**). *In vivo*, SsrBc resulted in lower SPI-2 induction and no increase in SPI-2 activity in acidic medium. Correspondingly, SsrBc did not support optimal SPI-2 activation and intracellular replication during HeLa cell infection (**Fig. 1f,g**). Therefore, although SsrBc is capable of DNA binding, it was insufficient to confer pH sensing or full activity during infection.

Interestingly, while we previously demonstrated a 5-fold acid-stimulated increase in the DNA binding affinity of SsrB to the ancestral *csgD* promoter (i.e. non-SPI-2); this work revealed a 32-fold increase in its affinity to the SPI-2 promoter *sseI*. This finding was also in agreement with previous AFM studies, which showed abundant binding of SsrB to the SPI-2 promoter at acid pH as compared to almost no binding observed at neutral pH (Liew et al., 2019). As the promoter recognition site of SsrB is a degenerate A-T rich 18 bp palindrome (Walthers et al., 2007; Tomljenovic-Berube et al., 2010), this increased affinity towards a SPI-2 promoter at acid pH is designed to selectively activate SPI-2 transcription within the macrophage vacuole during infection.

The non-conserved histidines His28, His34 and His72 are located in solvent-exposed regions in the predicted structure of SsrB (**Supplementary Fig. 2**). Their substitution did not have any effect on the acid-stimulated DNA binding of SsrB. Surprisingly, it was the conserved residue His12, which is solvent-accessible and in the vicinity of active site residues, that confers pH sensing to SsrB. In the i-TASSER and Alphafold predicted structures of SsrB (Jumper et al., 2021; Yang et al., 2015), His12 can form a H-bond with Asp11, it could also form a π-cation interaction with Lys106 (**Supplementary Fig. 5**). In a glutamate substitution (H12Q), the H-bond with Asp11 could potentially be maintained, and this could contribute to the full activity we observed at neutral pH (**Fig. 3b**). In the tyrosine/phenylalanine substitutions, the π-cation interaction with Lys106 would be maintained, and could potentially contribute to the acid pH-induced increase in SsrB activity (Nakajima et al., 2011). While the aromatic nature of the imidazole ring appears to contribute to pH sensing in SsrB, a role for His12 protonation at acid pH cannot be ruled out, as the basic amino acid substitutions lysine and arginine could possess reduced activity due to steric hindrance, making it difficult to assess the effect of protonation changes on activity. In summary, it appears that the mechanism behind SsrB pH sensing involves the aromatic nature of the imidazole side chain of His12, and this side chain may interact with Lys106 in the active site to influence activity. NMR structural analysis will hopefully reveal the interactions of His12 with other residues in the context of pH sensing.

As His12 is located near the SsrB active site, it is unsurprising that it influences SsrB phosphorylation. A similar phenotype was observed for the RR DegU, where an H12L substitution resulted in a lower rate of both phosphorylation and dephosphorylation, leading to a longer lifetime of DegU∼P (Dahl et al., 1992). An interesting finding was the elimination of pH dependence in SsrB phosphorylation upon substitution of His12. In the crystal structures of the NarL/FixJ subfamily RRs RcsB, VraR, and LiaR solved in the presence of the phosphomimetic BeF3, the conserved histidine residue is oriented in such a way that it can participate in a π-cation interaction with the Mg^2+^ ion essential for phosphorylation (Casino et al., 2018; Davlieva et al., 2016; Kumar et al., 2022; Leonard et al., 2013). In the unphosphorylated form, the histidine side chain of these same RRs faces away from the active site. Changes in the His12 rotameric state could thus modulate phosphorylation via its ability to coordinate the Mg^2+^ ion. Again, NMR experiments will hopefully shed light on the interaction between Mg^2+^ and His12.

The DNA binding activity of full-length SsrB is highly cooperative at the SPI-2 promoter *sseI*. Although SsrBc can form a dimer (Carroll et al., 2009), its Hill coefficient was substantially lower than the wild-type (**Fig. 1a-d**). This result highlights a heretofore unappreciated property of the SsrB receiver domain in the formation of higher order structures that are important for a robust level of transcription observed within the macrophage. Substitution of His12 also reduced cooperative binding at acid pH. In an AlphaFold structure of the SsrB dimer, residues Asp11, His12 and Lys106 are all present at the dimer interface (**Supplementary Fig. 7**). The presence of His12 at the dimer interface has also been observed in crystal structures of RcsB, VraR and LiaR (Casino et al., 2018; Huesa et al., 2021; Kumar et al., 2022; Leonard et al., 2013). Based on our binding data (**Fig. 3d,e**), and the structural models, His12 is required for SsrB oligomerization at acidic pH.

This work thus identifies the conserved His12 as playing a critical role in pH sensing, phosphorylation, and formation of higher order structures in SsrB and highlights its importance during *Salmonella* infection (**Fig. 6**). Whether or not this conserved histidine plays a similar role in other NarL/FixJ subfamily RRs has been unexplored to date, but would seem likely based on its high degree of conservation and the existing structural data. It was surprising that pH sensing appeared to be solely determined by a single amino acid in the phosphorylation domain of SsrB. Eliminating pH sensing via histidine 12 substitution rendered *Salmonella* completely avirulent and unable to survive and replicate in the vacuole (**Fig. 4**). Preventing pH switching as a means of controlling virulence is thus a novel strategy for controlling *Salmonella* pathogenesis.

**Fig. 6:**
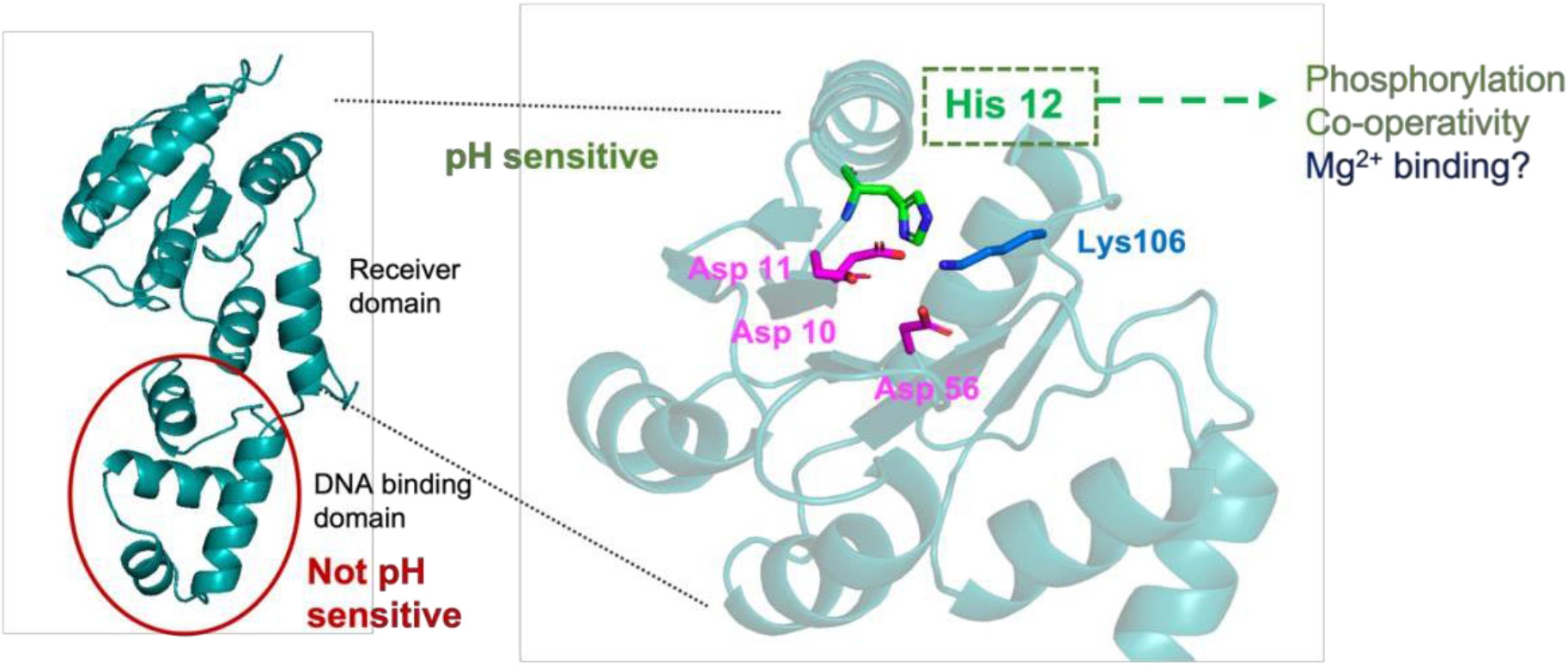
Summary of key findings. SsrB pH-sensing lies in the receiver domain and not in the DNA binding domain. A conserved Histidine at position 12, which lies in the proximity of active site residues, contributes to pH sensing. His12 also positively influences phosphorylation and cooperativity of DNA binding, and potentially influences Mg^2+^ binding as well.

## MATERIALS AND METHODS

### Strains and growth conditions

Bacterial strains and plasmids used in this study are listed in Methods Table 1. *Salmonella* Typhimurium (ST) strains were grown in either Luria Broth (Difco) or magnesium minimal medium (MGM): 100 mM Tris-HCl or MES-NaOH for pH 7.4 and 5.6, respectively, 5 mM KCl, 7.5 mM (NH_4_)_2_SO_4_, 0.5 mM K_2_SO_4_, 1 mM KH_2_PO_4_, 8 μM MgCl_2_, 0.2% glucose and 0.1% casamino acids (Beuzón et al., 1999). The bacteria were grown at 37°C shaking at 250 rpm. Transformation of plasmids carrying SsrBc or SsrB constructs into ST was performed using electroporation. HeLa cells used were maintained in Dulbecco’s modified Eagle medium (DMEM) containing sodium pyruvate, essential amino acids, 10% (v/v) fetal bovine serum (Gibco) and 1% (v/v) Penicillin/Streptomycin (P/S) at 37°C and 5% CO_2_.

### Molecular biology techniques, cloning and protein purification

DNA manipulations and cloning were performed using standard protocols described in Sambrook, 1989. Enzymes, reagents and DNA purification kits were purchased from ThermoFisher, New England Biolabs and Qiagen. Oligonucleotides were ordered from IDT Asia and are listed in Methods Table 2. SsrB histidine substitutions were generated by performing site-directed PCR mutagenesis using the pMPMA5Ω -SsrB wild-type plasmid (Feng et al., 2004) as the template. Sequences of the final constructs were verified by Axil Scientific, Singapore. Cloning and maintenance of plasmids used *E. coli* DH10B (TOP10). His_6_-tagged SsrB, SsrB H12Q and SsrBc were purified as previously described (Carroll et al., 2009).

### Single molecule unzipping assay of DNA binding

The DNA hairpin was prepared as previously described (Gulvady et al., 2018), using the 47 bp region of the *sseI* promoter which was protected from DNase I in the presence of SsrB (Feng et al., 2004) (Sequence in Methods Table 2). Preparation of the flow channel, attaching the DNA hairpin to the channel and the single molecule unzipping assay were performed as previously described (Gulvady et al., 2018; Liew et al., 2019). Briefly, in each experiment, the minimum hairpin unzipping force, known as the critical force, was determined in the absence of the protein. Binding events, represented by a delay in unzipping of the hairpin at the critical force, in presence of a given concentration of SsrB were measured over the course of 32 force cycles at pH 7.4 (50 mM KCl, 10 mM Tris) or 6.1 (50 mM KCl, 10 mM MES). The assay was then repeated with purified SsrB H12Q or SsrBc at pH 7.4 or pH 6.1. The probability of binding events was plotted as a function of the protein concentration, and the curve obtained by fitting the graph to the Hill equation using Origin was used to determine the *K*_*D*_. Three to five independent hairpins were assayed for each protein concentration.

### *β*-galactosidase assays

*β*-galactosidase assays were performed as described previously (Desai et al., 2016; Feng et al., 2003). DW637 strains carrying the pMPMA5Ω-SsrBc or SsrB constructs were grown overnight in LB containing 100 μg.ml^-1^ Ampicillin, sub-cultured 1:100 in 5 mL MGM medium pH 7.4 or pH 5.6. At early exponential phase (OD_600nm_ = 0.36-0.38, 3h 10 min), arabinose was added to a final concentration of 0.1% to the cultures. At 3 hours post induction, the OD of 200 μl of the diluted culture (1:2) was measured at 595 nm in a 96-well microtiter plate. 50 μl of this diluted culture was added to 450 μl lysis buffer containing 0.01% SDS, 50 mM *β*-mercaptoethanol and 20 μl CHCl_3_ in Z-buffer (Miller, 1972). The mixture was vortexed and incubated on a nutator for 5 min to lyse the cells. 100 μl of ONPG (ortho-nitrophenyl-*β*-galactoside, 4 mg.ml^-1^ in Z-buffer) was added to this mixture and incubated at 37ºC until a yellow-coloured product was formed. The reaction was stopped by adding 250 μl of 1M Na_2_CO_3_. 200 μl of the reaction mixture supernatant was transferred to a 96-well microtiter plate, and the OD was measured at 415 nm and 550 nm. The *β*-galactosidase activity was determined using the formula:

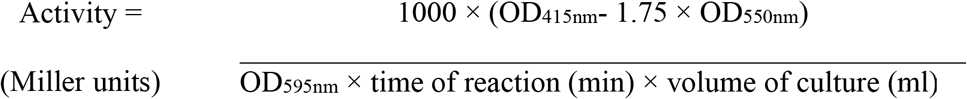

A minimum of three independent experiments were performed for each SsrB construct.

### Immunoblotting

DW637 strains carrying pMPMA5Ω, SsrB WT, H12Q or H12Y were grown in MGM pH 7.4 (10 ml each) and induced for the *β*-galactosidase assay as described above. Bacteria were harvested by centrifugation at 13000 rpm, washed once and resuspended in Tris buffer (pH 6.8), and sonicated (Fisher Scientific Sonic Dismembrator Model 100, setting 4, three pulses of 15 s). The lysed suspension was centrifuged at 13000 rpm, 4ºC for 20 min to separate the debris and the supernatant was used as the cell-free extract. The cell-free extract was mixed with 4X SDS-PAGE dye in a 3:1 ratio and heated at 95ºC for 10min. The samples were then separated on a gradient gel (4-20% Mini-PROTEAN® TGX™ Precast Protein Gels) with standard running buffer (0.4% (w/v) SDS, 25 mM Tris, and 192 mM glycine) at 125V. The separated samples were transferred to a PVDF membrane (Trans-Blot Turbo Mini 0.2 μm PVDF Transfer Packs) through semi-dry transfer (Bio-Rad Trans-Blot Turbo Transfer System). The PVDF membrane was blocked for 1 hour at room temperature with 5% Bovine serum albumin (BSA) in Tris-Buffered Saline containing 0.1% Triton-X-100 (TBS-T). The membrane was cut into two parts between the 35 and 55 kDa molecular weight markers. The lower molecular weight part of the membrane was incubated overnight at 4ºC in anti-6X-HisTag antibody (1:1000 dilution in blocking buffer, Invitrogen MA1-21315-HRP) to stain for SsrB protein constructs. The higher molecular weight part of the membrane was incubated overnight at 4ºC in anti-DNAK antibody (1:5000 dilution in blocking buffer, Abcam#69617 Mouse monoclonal [8E2/2]) as the loading control. Both membranes were given 5 washes with TBS-T at room temperature. The higher molecular weight membrane was incubated for 1 hour with HRP-tagged anti-mouse IgG (1:10000 dilution in 1% BSA in TBS-T, Santa Cruz Biotech goat anti-mouse IgG-HRP) at room temperature, followed by 3 washes with TBS-T. Both membranes were visualised using Clarity Max™ Western ECL Substrate (**Supplementary Fig. 4**).

### HeLa cell survival and *β*-galactosidase assays

HeLa cell infections were performed as described previously (Walthers et al., 2007). For the intracellular survival assay, 5×10^4^ HeLa cells were seeded and grown for 24 hours in 24-well plates, washed twice with Dulbecco’s Phosphate buffered saline (DPBS) and incubated in 0.5 ml DMEM without P/S containing 0.1% arabinose before infection. Overnight grown cultures of DW637 strains carrying pMPMA5Ω-SsrBc or SsrB constructs were diluted 1:30 in 3 ml LB and grown until late stationary phase. The bacteria were diluted to a final OD_600_ of 0.2 in DPBS. 83 μl of the diluted bacterial culture was added to the HeLa cells (multiplicity of infection = 200), and incubated for 30 min. To remove the unattached bacteria in the culture medium, the cells were washed twice with DPBS. Fresh DMEM containing 0.1% arabinose and 100 μg.ml^-1^ gentamicin was added (0 hour post infection, hpi) and the cells were incubated for 1 h. For the rest of the infection, the cells were incubated in DMEM containing 0.1% arabinose and 20 μg.ml^-1^ gentamicin. Intracellular ST were harvested at 2 and 16 hpi as follows: HeLa cells were washed with DPBS, and incubated in 0.5 ml of 0.1% Triton-X-100 in PBS for 10 min. After mixing by vigorous pipetting, 100 μl of the lysate was used to prepare serial dilutions in PBS for viable counting.

For the intracellular *β*-galactosidase assay, 2×10^5^ HeLa cells were seeded in 6-well plates and infection was performed as described above while increasing the volume proportionately. At 8 hpi, cells were washed once with DPBS and incubated in 1 ml of 0.1% Triton-X-100 in PBS for 10 min. The mixture was vigorously mixed and centrifuged at 500×*g* for 30 sec at 4ºC. The supernatant was centrifuged at 13000 rpm for 10 min, and washed once with PBS. The final pellet was resuspended in 120 μl of PBS, of which 10 μl was used to prepare serial dilutions for viable counting and 100 ml was used to perform the *β*-galactosidase assay. The final activity was measured as follows: Activity = (OD of the reaction mixture at 420nm × 10^9^) ÷ (Time of reaction in min × colony forming units.ml^-1^). A minimum of three independent experiments were performed for each SsrB construct.

### Phosphorylation assay

All phosphorylation assays were performed at room temperature using a phosphorylation buffer of pH 7.4 or pH 6.1, with phosphoramidate (PA) as the phosphodonor (Phosphoramidate preparation: (Stokes, 1893)). Purified SsrB WT or SsrB H12Q was added to a 100 μl reaction at a final concentration of 12 μM. The reaction mixture contained 1X phosphorylation buffer (50 mM HEPES-NaOH pH 7.4, or 50 mM MES-NaOH pH 6.1, 50 mM KCl, 20 mM MgCl_2_), with the appropriate amount of PA (0.5-100 mM final concentration). The reaction was immediately mixed after adding PA and incubated for the specified amount of time for the time-dependent reactions, or for 10 min for the concentration-dependent reactions. After incubation, the reaction was centrifuged for 1 min at 13000 rpm, and 75 μl of the supernatant was injected into an HPLC instrument (Waters 1525 binary pump) with a C_4_ reverse-phased column (Vydac 214TP54) and water:acetonitrile (containing 0.1% trifluoroacetic acid) solvent system. The following method was used: 40% acetonitrile (5 min), 40-50% acetonitrile gradient (20 min), 90% acetonitrile (7 min), 40% acetonitrile (8 min). The area under each observed peak was analysed with Breeze software. For the time-dependent reactions, the plot of the percentage of phosphorylated protein against time was fitted to a single exponential decay to obtain the *B*_*ma*x_ and *K*_*obs*_ using Prism software. The plot of the percentage of phosphorylated protein against the PA concentration was fitted to the Michaelis Menten equation using Prism to obtain the amount of PA required to phosphorylate 50% of the protein (*K*_*M*_) and the *V*_*max*_ of the reaction. The *K*_*cat*_ was obtained by dividing the *V*_*max*_ by total SsrB concentration (12 μM), and phosphorylation efficiency was calculated by dividing *K*_*cat*_ by *K*_*M*_.

**Methods Table 1:**
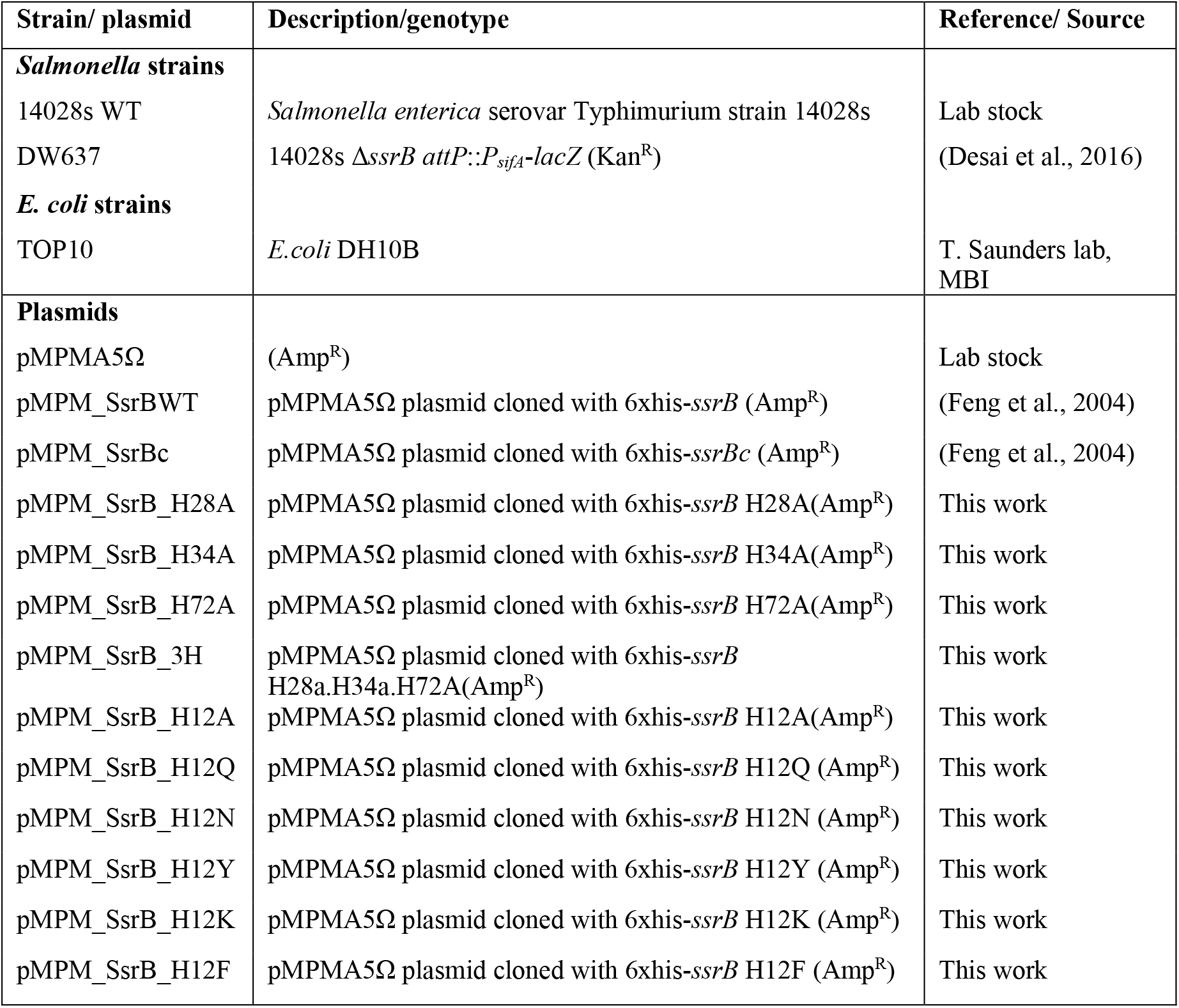
Strains and plasmids used in this study

**Methods Table 2:**
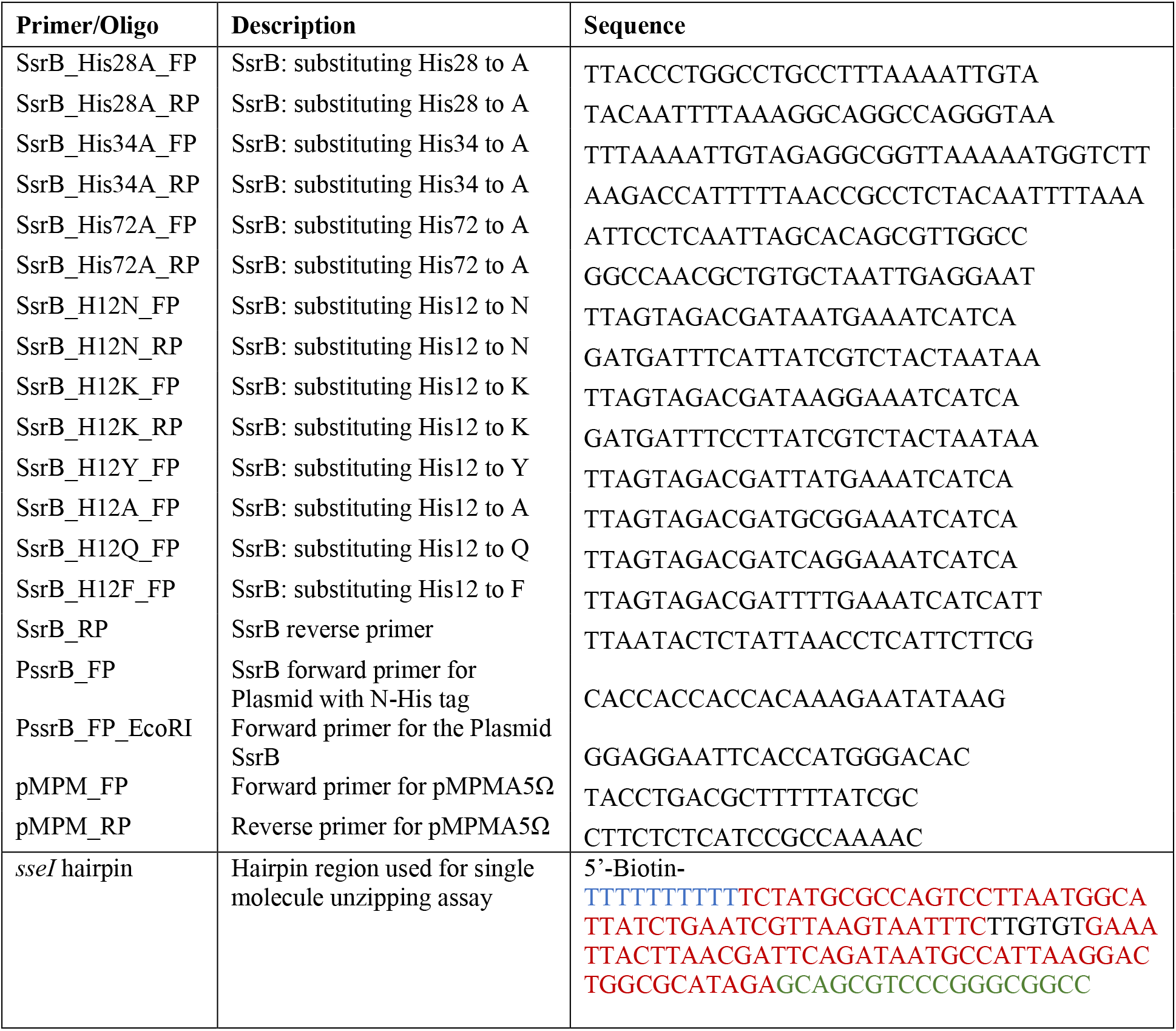
Primers and oligonucleotides used in this study

## Abbreviations

SPI-2: *Salmonella* pathogenicity island 2
SCV: *Salmonella*-containing vacuole
bp: base pairs
RR: Response regulator
TCRS: Two component regulatory system
SK: sensor kinase
ST: *Salmonella* Typhimurium
*K*_*D*_: Dissociation constant
Spt-PALM: Single particle tracking Photoactivated localization microscopy
hpi: hours post infection
c.f.u: colony forming unit

## ACKNOWLEDGEMENTS

We are grateful to Prof. Yan Jie (MBI, NUS, Singapore) for providing laboratory space and Dr. Ranjit Gulvady for advice and instruction on the single molecule experiments. This work was supported in part by a Regional Centre of Excellence Grant from the Ministry of Education to the Mechanobiology Institute, National University of Singapore. LJK and DS were also supported by start-up funds from UTMB, a Texas STAR award and CPRIT RP200650 to LJK.

