## Supplementary info for "A pH-sensitive switch activates virulence in *Salmonella*"

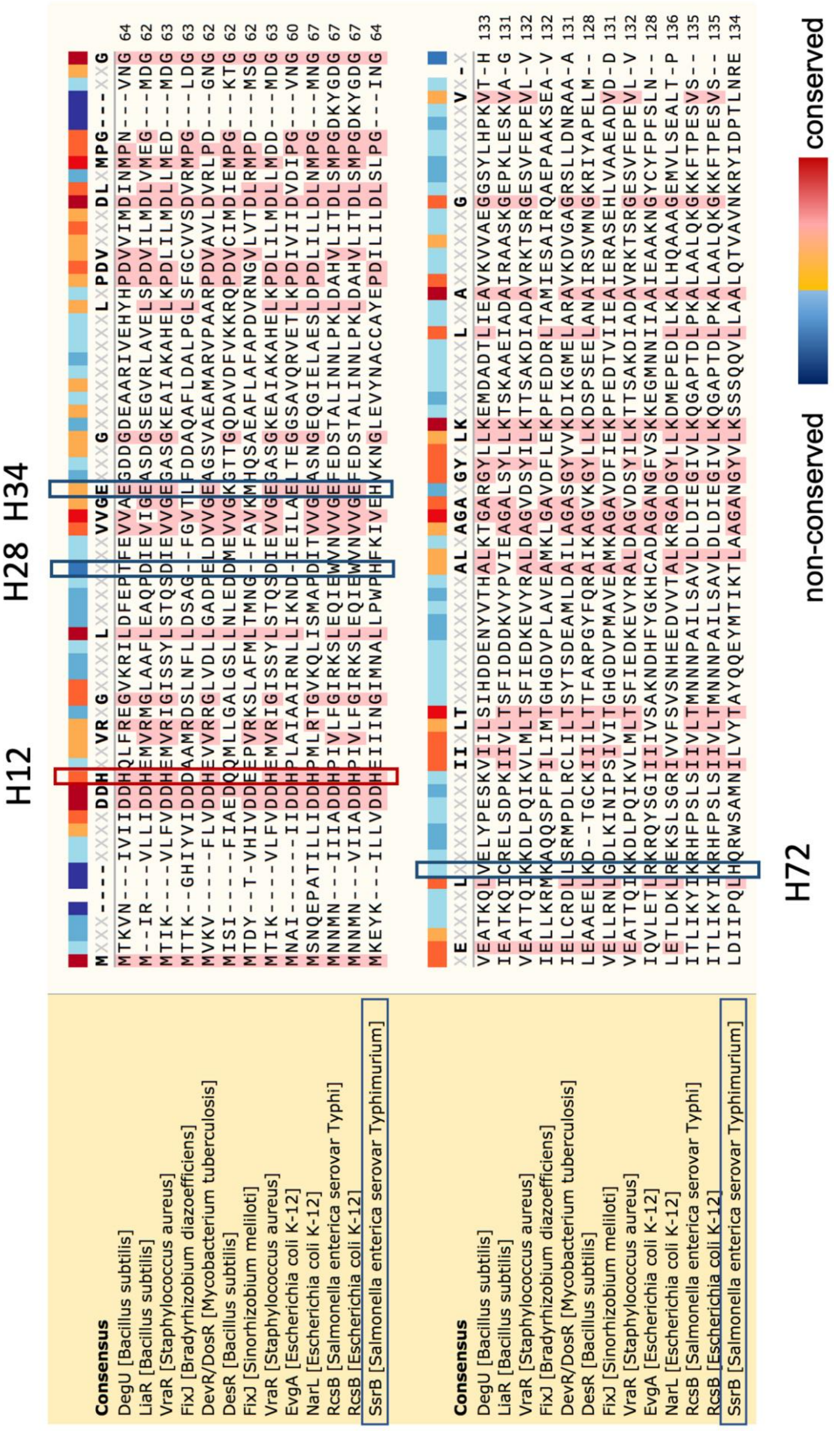

**Supplementary Fig. 1: Analysis of NarL/FixJ RR receiver domains and selection of histidine residues.** Multiple sequence alignment of the receiver domains of NarL/FixJ sub-family RRs from sequences obtained through a DELTA_BLAST of SsrB sequence against the UniProt/SWISSPROT database. These sequences share >20% identity with SsrB. The histidines at positions 28, 34 and 72 in SsrB were the least conserved amongst NarL/FixJ family members, and their homologous aligned residues are shown in the blue box. The histidine at position 12 in SsrB was highly conserved among most RRs, and its homologous aligned residues are shown in the red box. Amino acid conservation is represented by the colored bar above the aligned residues, with blue being the least conserved and red being the most conserved residues.

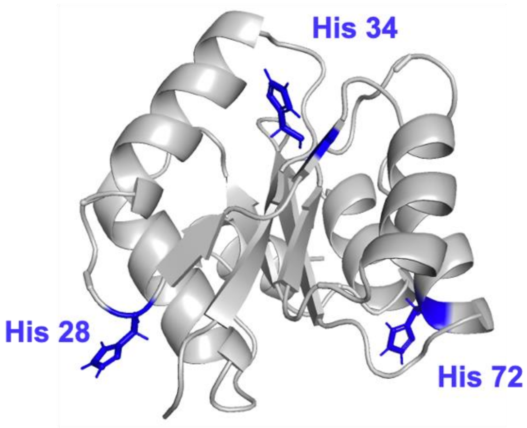


**Supplementary Fig. 2:** The location of His28, His34 and His72 on the predicted receiver domain of SsrB (visualized using PyMol). His28 is present in the loop between $\alpha 1$ and $\beta 2$, His34 is on $\beta 3$ and His72 is on $\alpha 3$.

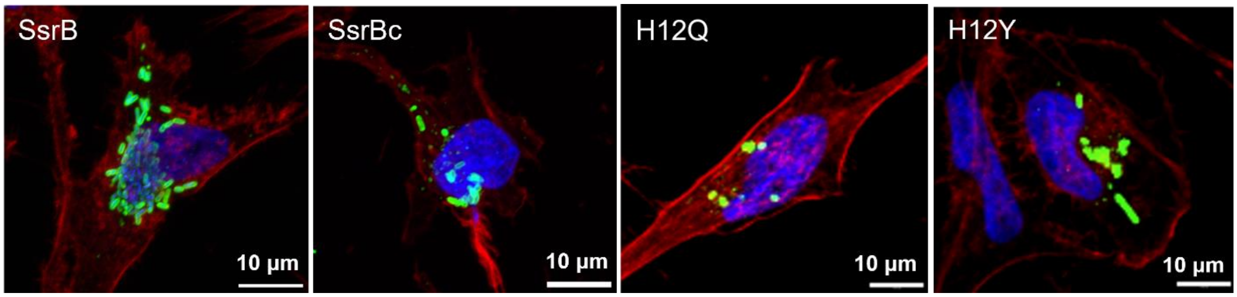


**Supplementary Fig. 3:** DW637 strains expressing SsrB, SsrBc, H12Q or H12Y infecting HeLa cells at 16 hpi. *Salmonella* were stained with anti-LPS antibody (green), HeLa cells were stained using phalloidin (red) and DAPI (blue).

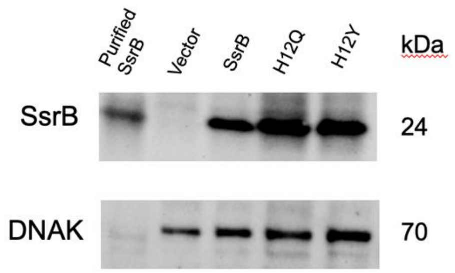


**Supplementary Fig. 4:** Immunoblotting of strains expressing various SsrB constructs grown in MGM pH 7.4. An anti-6X-HisTag monoclonal antibody was used to detect SsrB, and an anti-DnaK antibody was used as a loading control. Lane 1 contains purified SsrB (6.9 µg) as a positive control. Lanes 2 to 5 contain cell-free extracts from strains expressing the empty vector (lane 2), SsrB (lane 3), H12Q (lane 4), H12Y (lane 5).

**a.** **b.**

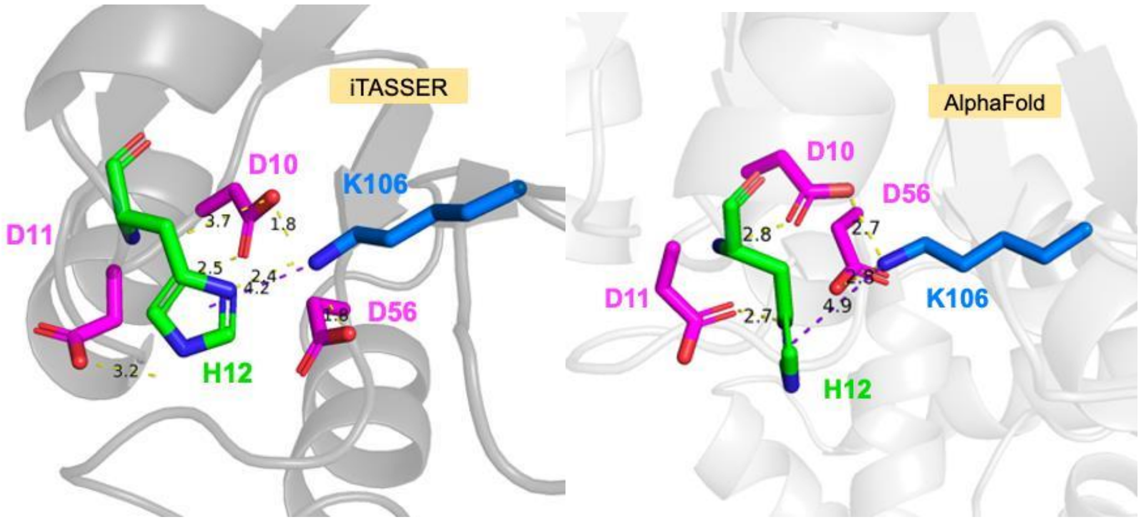


**Supplementary Fig. 5: His12 interactions with residues in the SsrB receiver domain.** A) In an i-TASSER prediction, His12 forms polar contacts with Asp10, Asp11 and Lys106, and a π-cation interaction with Lys106. B) In an AlphaFold prediction, His12 forms polar contacts with Asp10 and Asp11, and a π-cation interaction with Lys106. Structures were visualised using PyMol; yellow dashed lines represent polar contacts, and purple lines represent π-cation distance.

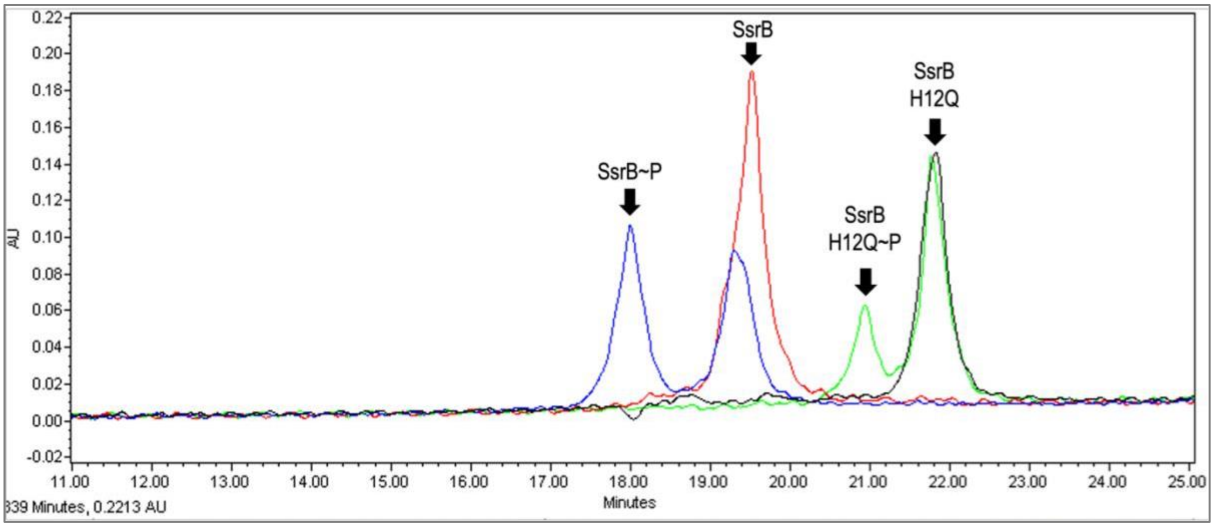


**Supplementary Fig. 6:** The elution profile of unphosphorylated and phosphorylated SsrB and SsrB H12Q in a 40-50% acetonitrile in water gradient. The red curve represents the profile of unphosphorylated SsrB, which has a retention time of 19.5 min. The blue curve represents a reaction of SsrB with 2.5 mM PA, with SsrB~P eluting at 18 min and SsrB eluting at 19.3 min. The black curve represents the profile of unphosphorylated SsrB H12Q, its retention time was 21.8 min. The green curve represents a reaction of SsrB H12Q with 2.5 mM PA, with SsrB H12Q~P eluting at 20.9 min and SsrB H12Q eluting at 21.8 min.

**a.** **b.**

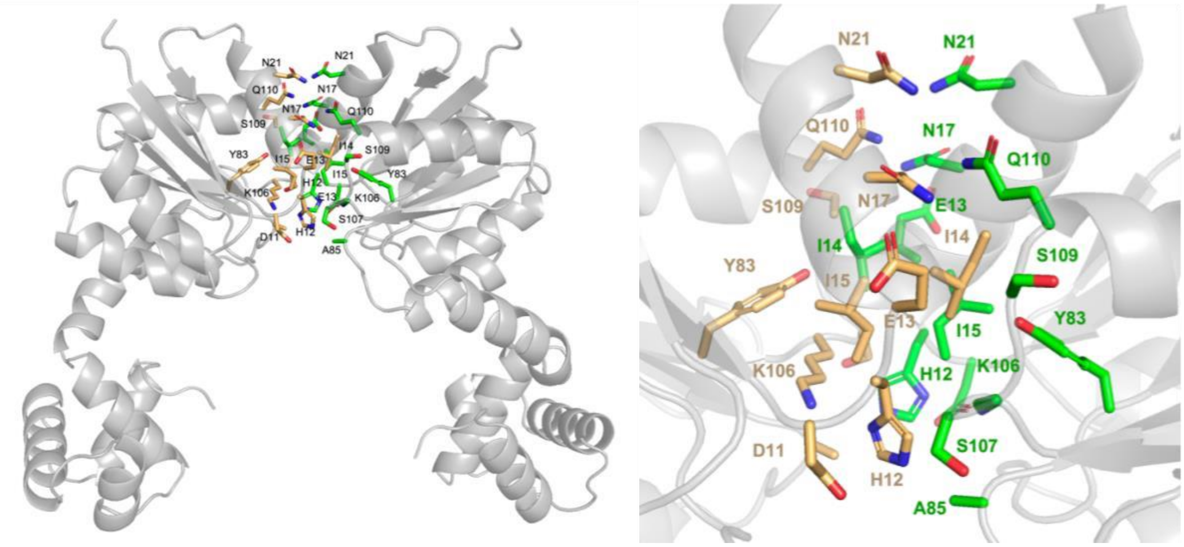


**Supplementary Fig. 7:** AlphaFold prediction of an SsrB dimer. A) The dimer structure of SsrB was predicted using the AlphaFold server. Interacting residues were analyzed using UCSF ChimeraX. Side chains of residues involved in the dimer interface are coloured and represented as sticks, with orange sticks representing residues from monomer one and green sticks representing residues from the other monomer. B) A zoomed in image of the dimerization interface.
